# Elucidation of Master Allostery Essential for Circadian Clock Oscillation in Cyanobacteria

**DOI:** 10.1101/2021.08.30.457330

**Authors:** Y. Furuike, A. Mukaiyama, D. Ouyang, K. Ito-Miwa, D. Simon, E. Yamashita, T. Kondo, S. Akiyama

**Affiliations:** Research Center of Integrative Molecular Systems (CIMoS), Institute for Molecular Science, National Institute of Natural Sciences, 38 Nishigo-Naka, Myodaiji, Okazaki 444-8585, Japan; Department of Functional Molecular Science, SOKENDAI (The Graduate University for Advanced Studies), 38 Nishigo-Naka, Myodaiji, Okazaki 444-8585, Japan; Division of Biological Science, Graduate School of Science and Institute for Advanced Studies, Nagoya University, 464-8602 Nagoya, Japan; Institute for Protein Research, Osaka University, 3-2 Yamada-oka, Suita 565-0871, Japan

## Abstract

Spatio-temporal allostery is the source of complex but ordered biological phenomena. To identify the structural basis for allostery that drives the cyanobacterial circadian clock, we crystallized the clock protein KaiC in four distinct states, which cover a whole cycle of phosphor–transfer events at Ser431 and Thr432. The minimal set of allosteric events required for oscillatory nature is a bidirectional coupling between the coil-to-helix transition of the Ser431-dependent phospho-switch in the C-terminal domain of KaiC and ADP release from its N-terminal domain during ATPase cycle. An engineered KaiC–protein oscillator consisting of a minimal set of the identified master allosteric events exhibited mono-phosphorylation cycle of Ser431 with a temperature-compensated circadian period, providing design principles for simple post-translational biochemical circadian oscillators.

**One Sentence Summary:** Coupling between a phospho-switch and KaiC ATPase-dependent nucleotide exchange drives the cyanobacterial circadian clock.

## Main Text

The cooperative nature of allosteric regulation is a source of nonlinearity that gives rise to oscillatory phenomena in diverse cellular functions (*1, 2*). Circadian clock systems are a typical example, wherein clock proteins are post-translationally phosphorylated at multiple sites in programmed (*3-5*) or pseudo-random manners (*6, 7*) as a means to allosterically regulate the stability of hetero-multimeric complexes of clock proteins (*8, 9*), delays for feedback loops (*6*), and period length (*10*). Accordingly, a great deal of effort has been devoted to characterizing the phosphorylation-dependent allosteric structural changes in the clock proteins along the circadian reaction coordinate (*8, 9, 11-15*).

The cyanobacterium *Synechococcus elongatus* PCC7942 is one of the simplest prokaryotic model organisms (*16*). Only two adjacent amino acid residues, S431 and T432, in the core clock protein KaiC are phosphorylation targets (*17, 18*). In the presence of KaiA, KaiB, and ATP, KaiC exhibits a phosphorylation cycle *in vitro* (P-cycle): ST → SpT → pSpT → pST → ST, where S, T, pS, and pT indicate S431, T432, phosphorylated S431 (pS431), and phosphorylated T432 (pT432), respectively (*17-19*). Whereas the KaiC P-cycle has been investigated *in vivo* (*20*), *in vitro* (*19, 21*), and *in silico* (*18, 22-24*), little is known about the allosteric regulation of the KaiC P-cycle because all KaiC structures reported to date share nearly identical conformations at the phosphorylation sites, irrespective of the presence or absence of phosphoryl modifications (*8, 9, 14, 15*).

To visualize the structural basis for allosteric oscillatory regulation, we crystallized the KaiC hexamer in eight distinct states and sorted them from the fully phosphorylated KaiC-pSpT to the fully dephosphorylated KaiC-ST so that fractional changes in pS431 and pT432 per hexamer in the crystalline phase (**Fig. 1A**) reproduce those observed during the circadian cycle in solution (**Fig. 1B**). Each subunit of the KaiC hexamer consists of an N-terminal ATPase (C1) domain (*11, 25*) and a C-terminal auto-kinase/phosphatase (C2) domain (*14, 21*) (left panel of **Fig. 1C**). Dephosphorylation of pT432 in the fully KaiC-pSpT hexamer occurred in a stepwise manner, but no specific order was observed in regard to which subunit was dephosphorylated at each step (fig. S1). However, systematic ADP accumulation was observed for the C1 domain (C1-ADP) of KaiC-pST (**Fig. 1, A and C**, and fig. S1). The accumulated C1-ADP molecules were replaced with C1-ATP during the transition from KaiC-pST to KaiC-ST (**Fig. 1C**). These results indicate that the auto-dephosphorylation events in the C2 domain are linked to C1-ATP hydrolysis (ADP production) and C1-ADP exchange.

**Fig. 1.**
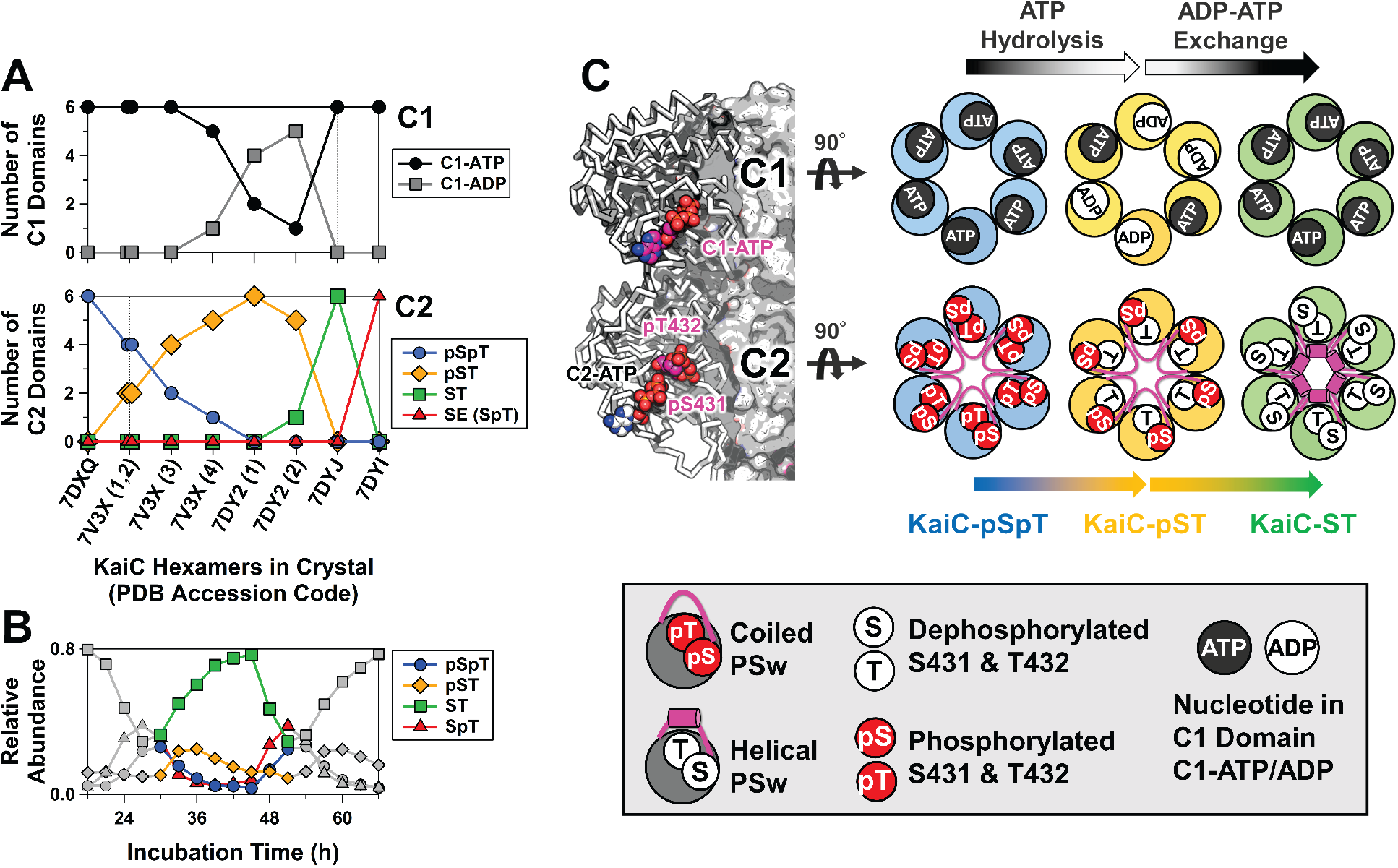
KaiC crystal structures sorted by phosphorylation, ATP hydrolysis, and nucleotide-exchange events. (**A**) Fractional changes for ATP/ADP bound in the C1 domain (upper panel) and phosphorylation status of S431 and T432 in the C2 domain (lower panel) in the crystalline phase. Horizontal axis depicts the PDB accession codes for nine KaiC hexamers crystallized in eight distinct states. For 7V3X and 7DY2, four and two hexamers present in each asymmetric unit are distinguished by the number in parentheses. Vertical axes represent the number of C1 and C2 domains that are in the ATP/ADP–bound and phosphorylated/dephosphorylated states, respectively, in the hexamer of interest. (**B**) Phosphorylation cycle for KaiC in solution. (**C**) Schematic drawings for three representative hexamers for KaiC-pSpT (7DXQ), KaiC-pST (7DY2), and KaiC-ST (7DYJ). Arrows above and below the hexameric rings represent biochemical reactions and state transitions, respectively, that are suggested by the sorted structures. The nomenclatures used in the drawings are defined in the bottom panel.

Our crystal-structure library indicates that key allosteric communication occurs between the C1 and C2 domains during the transition from KaiC-pST to KaiC-ST. We observed that the region upstream of S431 (T416–H429), which adopts a coiled structure in KaiC-pST, folds into a novel helical structure in KaiC-ST (left panels of **Fig. 2**). A helix-to-coil reversal was observed in the transition from KaiC-SpT to KaiC-pSpT (right panels of **Fig. 2**). These results enable us to assign the upstream region of S431 as a local phospho-switch (PSw). As shown in the left panels of **Fig. 2**, removal of the phosphoryl group from pS431 disrupted hydrogen bond interactions between pS431 and T426 in KaiC-pST, which provided the space necessary for the PSw to fold into the compact helical structure in KaiC-ST.

**Fig. 2.**
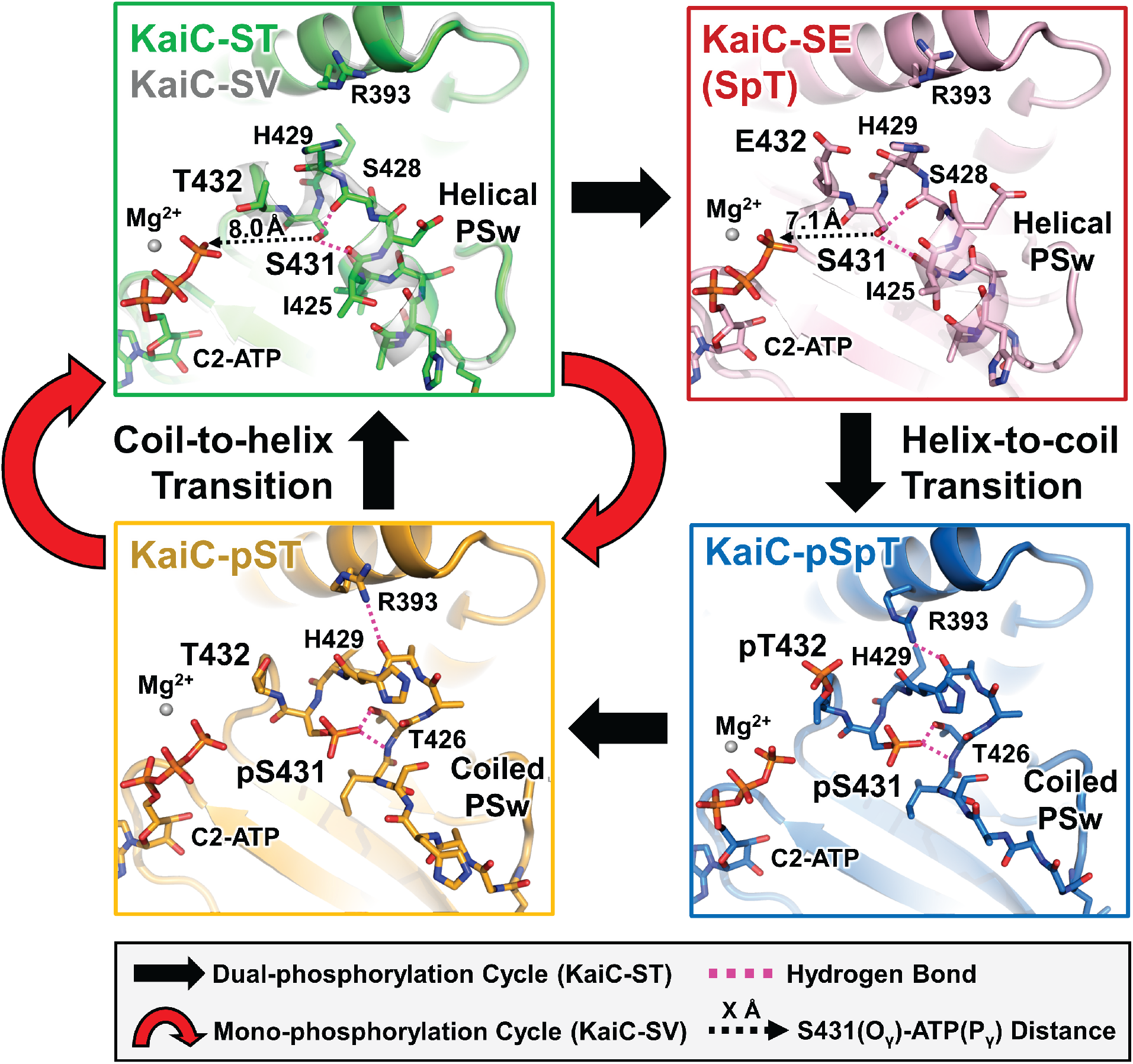
Cyclic structural changes in the phosphor-switch (PSw) located upstream of S431 in KaiC-ST (green), KaiC-SV (white), KaiC-SE (pink), KaiC-pSpT (blue), and KaiC-pST (orange). Black and red arrows represent the dual- and mono-phosphorylation cycles in KaiC-ST and KaiC-SV, respectively. Dashed magenta lines correspond to hydrogen bonds around S431/pS431. Dashed black arrows indicate the distances between the S431 O_γ_ atom and C2-ATP P_γ_ atom.

The PSw coil-to-helix transition in the C2 domain was coupled with the ADP release from the C1 domain. Indicated by arrows in **Fig. 3A** (gradient from orange to green), the ADP-to-ATP exchange at the C1–C1 interface resulted in a slight but systematic positional shift of the neighboring C1 domain. This rearrangement drove repositioning of the side chain of the basic residue R217 away from the neutral residue Q394 located in the C1–C2 interface (box in **Fig. 3A**), eventually disrupting the hydrogen bonding interaction between them that was observed in KaiC-pST. Instead, the acidic residue E214 captured the released Q394 through a new hydrogen bond, thereby causing the C2 domain to move closer to the C1 domain in KaiC-ST. R393 and H429 are particular examples of this C1-directed movement by up to ∼3 Å, assisting the coil-to-helix switching of the PSw through multiple hydrogen-bond formation of S431 with I425 and S428 (upper left panel of **Fig. 2**).

**Fig. 3.**
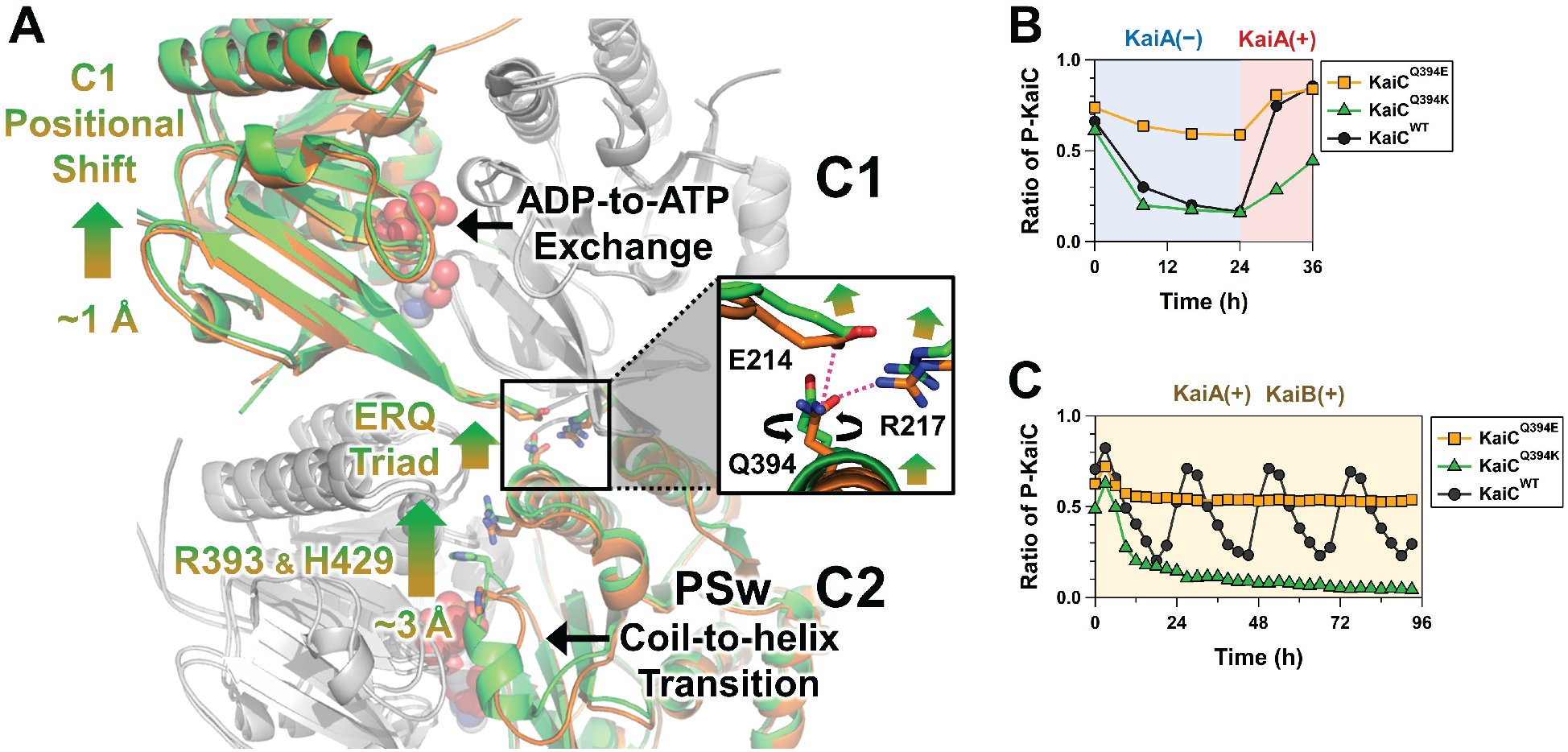
Structural basis for master allostery between C1-ATPase and C2-PSw in KaiC. (**A**) Tertiary and quaternary structural changes in the transition from KaiC-pST (orange) to KaiC-ST (green). The ADP–ATP exchange occurring at the C1–C1 interfaces and the coil-to-helix transition for PSw are allosterically coupled via rearrangements of E214, R217, Q394, R393, and H429. The inserted box indicates the zoomed-in-view of the ERQ triad mediating communication between the C1 and C2 domains. (**B**) Effects of Q394E and Q394K mutations on the switching ability of C1–C2 allostery. Proportion of phosphorylated KaiC (P-KaiC) was measured for 24 h at 30°C and after the KaiA addition at 24 h. (**C**) Effects of Q394E and Q394K mutations on the P-cycle.

The neutral character of residue 394 was critical to this allosteric communication because a Q394E mutation (KaiC^Q394E^) stabilized its electrostatic interaction with R217 but destabilized with E214, which resulted in constitutive phosphorylation, even with KaiC alone (**Fig. 3B**), and arrhythmic accumulation of its phosphorylated forms even in the presence of KaiA and KaiB (**Fig. 3C**). Consistent with this, KaiC^Q394K^ exhibited very slow phosphorylation rates in the presence of KaiA (**Fig. 3B**) and remained dephosphorylated even in the presence of KaiA and KaiB (**Fig. 3C**). Hence, most of the large-scale bidirectional C1–C2 communication during the P-cycle takes place through an E214–R217–Q394 (ERQ) triad in the transition from KaiC-pST to KaiC-ST.

Our biochemical analysis demonstrated that the observed C1–C2 communication is necessary and sufficient to allosterically drive the P-cycle. We designed a KaiC T432V mutant (KaiC-SV) to investigate the mono-phosphorylation/dephosphorylation of S431 with minimal structural perturbation around residue 432. Valine is closest to threonine in terms of topology and volume (fig. S2, supplementary text), as evidenced by conservation of the helical PSw in KaiC-SV (upper left panel of **Fig. 2**). Astonishingly, KaiC-SV exhibited an *in vitro* mono–P-cycle for S431 (**Fig. 4A**) (KaiC-SV ↔ KaiC-pSV, red arrows in **Fig. 2**), an *in vitro* ATPase cycle (**Fig. 4B**), and an *in vitro* assembly and disassembly cycle (fig. S3, supplementary text) with a prolonged but temperature-compensated period (51 ± 2 h, *Q*_10_ = 1.1) (**Fig. 4C**). By contrast, KaiC-CT, which was designed to allow for mono-phosphorylation/dephosphorylation of T432 with minimal structural perturbation around residue 431, was arhythmic at 30°C (**Fig. 4A**). Furthermore, KaiC-SE, which was generated to allow for mono-phosphorylation/dephosphorylation of S431 without going through the fully-dephosphorylated KaiC-ST, was also arhythmic (**Fig. 4A**). The minimal set of allostery that ensures oscillatory nature is the coupling between the PSw transition associated with mono-phosphorylation/dephosphorylation of S431 and nucleotide exchange in the C1 domain (left panels of **Fig. 2**).

**Fig. 4.**
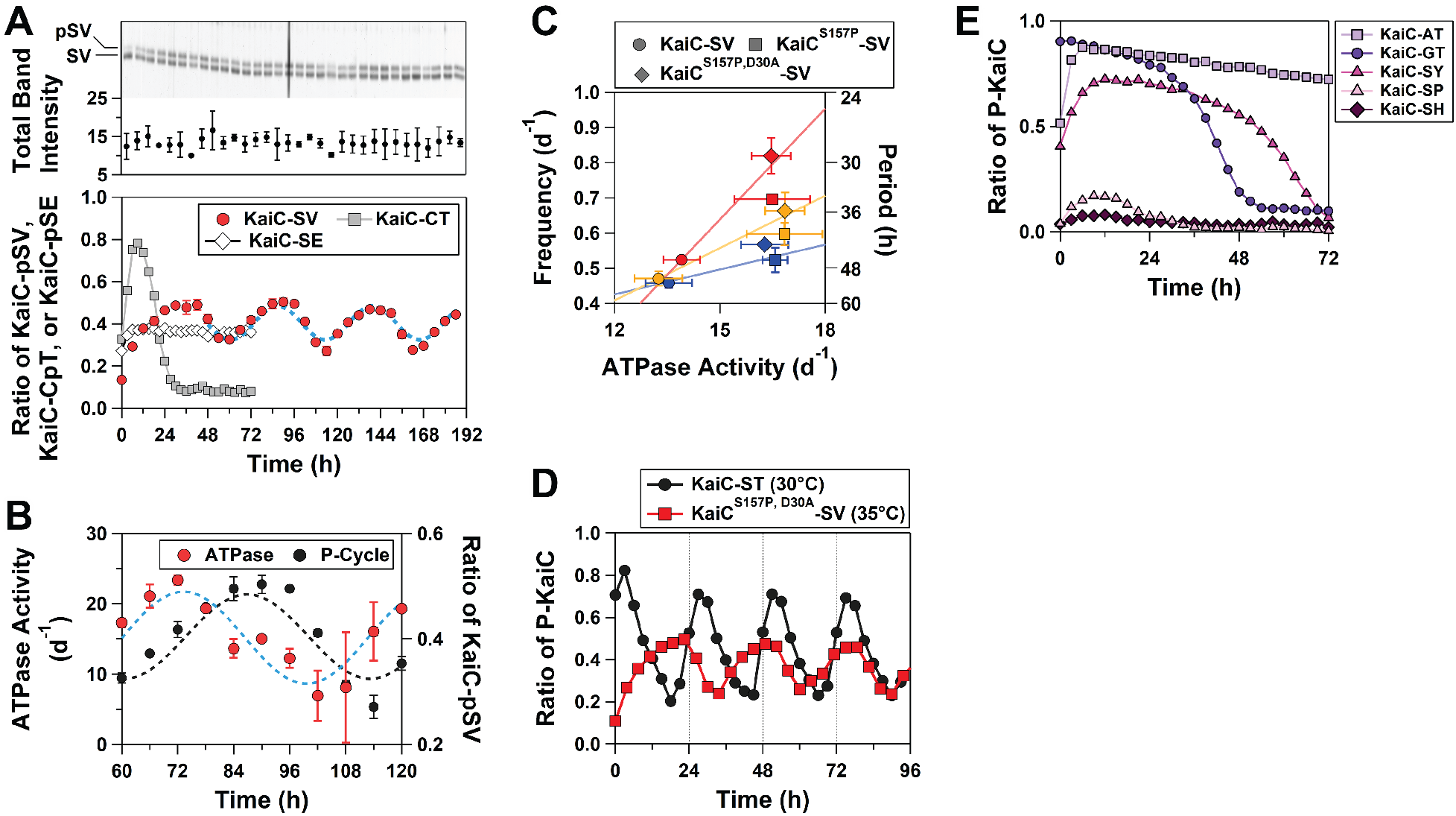
Mono-phosphorylation rhythms for KaiC-SV. (**A**) *In vitro* P-cycle at 30°C. Dotted line represents fitting of a cosine function to the data. Upper panel represents a magnified view of the upper phosphorylated (pSV) and lower dephosphorylated (SV) bands in SDS-PAGE gels. (**B**) *In vitro* ATPase rhythm at 30°C. (**C**) Correlation plot between ATPase activity for KaiC alone and the frequency (24/period) of the *in vitro* P-cycle in the presence of KaiA and KaiB. Blue, orange, and red markers correspond to data analyzed at 25, 30, and 35°C, respectively. (**D**) Near circadian mono–P-cycle (29 ± 2 h) for KaiC^S157P,D30A^-SV at 35°C. (**E**) Arrhythmicity of KaiC-AT, KaiC-GT, KaiC-SY, KaiC-SP, and KaiC-SH at 30°C.

The prolonged mono–P-cycle observed for KaiC-SV implies that phosphorylated T432 (pT432) plays a role in accelerating the KaiC-ST P-cycle. In fact, the O_γ_ atom of S431 was positioned 1 Å closer to C2-ATP in the KaiC-SpT–mimicking mutant KaiC-SE (upper right panel of **Fig. 2A**) relative to the position in KaiC-ST, which likely facilitates the subsequent formation of KaiC-pSpT. To investigate whether other mutations can allosterically suppress the period-prolonging T432V substitution, we introduced various ATPase-activating mutations into KaiC-SV. A S157P substitution (KaiC^S157P^-SV), which is a known period-shortening and ATPase-activating mutation (*11, 25, 26*), resulted in a less prolonged period (40 ± 2h) at 30°C (**Fig. 4C**). We also introduced the ATPase-activating mutation D30A (*27*). The resultant KaiC^S157P,D30A^-SV exhibited a near circadian mono–P-cycle (29 ± 2 h, **Fig. 4D**) at 35°C with a slight impairment in temperature compensation (*Q*_10_ = 1.4). However, it should be noted that the ATPase activities of KaiC^S157P^-SV and KaiC^S157P,D30A^-SV are temperature-compensated, as was observed for KaiC-SV (**Fig. 4C**). Combined with KaiA and KaiB, KaiC^S157P,D30A^-SV constitutes a simple biochemical post-translational circadian oscillator with a sole-phosphorylation site.

The crucial role of the observed allostery is further supported by evolutional conservation and divergence of the dual phosphorylation sites. Multiple sequence alignment of KaiC and its homologues from other organisms revealed a near-perfect conservation of S431 (*28*). Only alanine (*Fibrella aestuarina* BUZ 2) and glycine (*Allochromatium vinosum* DSM 180) were observed exceptions, but the arrhythmicity of KaiC-AT and KaiC-GT in the presence of KaiA and KaiB supports our present interpretation (**Fig. 4E**). By contrast, amino acids other than threonine, such tyrosine (*Pseudomonas oryzihabitans*), histidine (*Massilia sp*. WG5), and proline (*Pontibacter korlensis*), are observed at position 432. We generated KaiC-SY, KaiC-SH, and KaiC-SP to investigate the effects of these substitutions and found that all of them exhibited arrhythmicity (**Fig**.**4E**). These results suggest that different evolutionary selection pressures acted on the two sites; position 431 was conserved to allow for acquisition of master C1–C2 allostery that gives rise to the temperature-compensated rhythmicity, and amino acid variances at position 432 fine-tune the period length into the circadian time scale by accelerating the C1–C2 allostery mediated by S431.

Mathematical modeling approaches have highlighted the importance of multisite phosphorylation in oscillatory phenomena. Simple enzymatic futile cycles in which a substrate shuttles between two states, e.g., mono-phosphorylated and dephosphorylated forms, via forward and backward reactions catalyzed by an opposing enzyme converge into an equilibrium (*29*). Unbiased systems targeting a substrate with two modification sites retain a limited (∼0.1%) potential to exhibit self-sustained oscillation (*30*). This apparent inconsistency can be resolved by taking into consideration the fact that KaiC-SV can still be regarded as a two-site system because its single site for phosphorylation is influenced allosterically by another biochemically active site for ATP hydrolysis in the C1 domain (**Fig. 3A**).

The cyanobacterial circadian clock is the simplest circadian clock system known to date in terms of the number of components, but mechanistically it is a very complex system involving the ATPase cycle and a four-state P-cycle. The oscillators we designed using KaiC-SV and KaiC^S157P,D30A^-SV reconstruct the complex system into a minimal unit by extracting the core allostery from the complex cycle. This will serve as a research tool for further elucidation of mechanisms, such as period determination, temperature compensability, and entrainment, and will provide design principles for the simplest post-translational biochemical oscillator that oscillate with a temperature-compensated circadian period.

## Acknowledgments

Diffraction data were collected at BL44XU at the SPring-8 facility under the proposals 2017A6700, 2017B6700, 2018A6700, 2018B6700, 2019A6700, 2019B6700, 2020A6700, 2020A6500, 2017A6702, 2017B6702, 2018A6802, 2018B6802, 2019A6902, 2019B6902, and 2020A6502. This research was partly supported by the Platform Project for Supporting Drug Discovery and Life Science Research (BINDS) from AMED under Grant Number JP20am0101072.

## Author contributions

Y.F., A.M., and S.A. designed the experiments. Y.F. collected the diffraction data and analyzed it with input from E.Y. Y.F., A.M., D.O., and D.S. performed the biochemical experiments. K.I.-M. and Y.F. performed *in vivo* experiments with input from T.K. S.A. and Y.F. drafted the manuscript with input from all authors.

## Data and materials availability

The atomic coordinates and structure factors are deposited in the Protein Data Bank with accession codes 7DYK (KaiC-SV), 7DYJ (KaiC-ST), 7DY2 (KaiC-pST), 7DYI (KaiC-SE), 7DXQ (KaiC-pSpT), and 7V3X (KaiC-pSpT & pST). Requests for data or materials should be addressed to.

## Supplementary Materials

Materials and Methods

Figures S1-S3

Tables S1 References (*31-40*)

