## supplementary for "Elucidation of Master Allostery Essential for Circadian Clock Oscillation in Cyanobacteria"

**This PDF file includes:**

Materials and Methods
Supplementary Text
Figs. S1 to S3
Tables S1
References (*31-40*)

### Materials and Methods

Expression and purification of KaiC

Plasmid vectors for wild-type KaiC and KaiC mutants were generated for glutathione S-transferase (GST)-tagged (pGEX-6P-1) or hexa-histidine (His)-tagged (pET-3a) proteins. Kai proteins were expressed in BL21(DE3) or BL21(DE3)pLysE *E. coli.* cells and purified (*17, 27, 31*).

*In vivo* and *in vitro* rhythm assays

*In vivo* bioluminescence assays were conducted as previously reported using a cooled-CCD camera system (*32*). KaiC phosphorylation and ATPase cycles were reconstituted *in vitro* in the presence of KaiA, KaiB, and ATP (*19*). The relative abundances of the four phosphorylation states of KaiC were quantified using SDS-PAGE and the LOUPE software (*33*). The ATPase activity of KaiC was measured as previously described (*13, 27*) and presented as the number of ATP molecules hydrolyzed into ADP molecules per KaiC monomer per unit time. Unless otherwise noted, all the rhythm assays were conducted at 30°C.

Crystallization of KaiC

All crystals were obtained using the vapor diffusion method. The purified samples were concentrated to 3.5 mg/mL and stored in a solution of 20 mM Tris-HCl (pH 8.0), 150 mM NaCl, 5 mM $MgCl_2$, 1 mM DTT, and 1 mM ATP. Crystals for KaiC-pSpT and KaiC-pST were obtained in solutions containing 80–250 mM acetic acid and 1.0–1.5 M sodium acetate and were frozen with 30% (w/v) glycerol or 25% (w/v) PEG8000. Crystals that formed in the $P6_3$ space group were prepared in solutions containing 100 mM Tris-HCl (pH 7.0), 1 M KCl, 0.7–1.0 M sodium/potassium tartrate, 0.3–1.2 M sodium acetate, and 1–5 mM AMP-PNP.

Data collection and structure determination

X-ray diffraction data were collected on beamline BL44XU at SPring-8 (Harima, Japan). Crystals were mounted at 100 K under a cryostream, and diffractions images were recorded with an MX-225 (Rayonix) or PILATUS (Eiger) detector and processed using HKL2000 (*34*) and XDSGUI (*35*).

Initial phases were obtained using molecular replacement with previously deposited structures (2GBL (*36*) or 7DYJ) and MOLREP (*37*). Refinement and modeling were conducted using Refmac5 (*38*) and COOT (*39*), respectively. The crystal structure of KaiC-ST was refined using reflections merged from two crystals that had only a 0.7% difference in lattice sizes ($a_{\mathrm{crystal1}}$ = 94.7 Å and $c_{\mathrm{crystal1}}$ = 180.5 Å; and $a_{\mathrm{crystal2}}$ = 94.6 Å and $c_{\mathrm{crystal2}}$ = 181.8 Å) (table S1). The asymmetric unit for the $P6_3$ crystal contained a dimer of KaiC-ST and was arranged along a crystallographic three-fold axis to form a hexamer. The KaiC-SE and KaiC-SV crystals also belonged to the $P6_3$ space group. KaiC-pSpT crystallized in the space group $P2_12_12_1$ with a hexamer in the asymmetric unit, whereas KaiC-pST crystallized in the space group $P2_1$ with two hexamers in the asymmetric unit. Graphic representations were generated using PyMOL (Schrödinger). The statistics for data collection and refinement are listed in table S1.

KaiA–KaiB–KaiC interaction assay

KaiA (0.04 mg/mL) and/or KaiB (0.04 mg/mL) were incubated with KaiC-SV (0.2 mg/ml) at 30°C in a buffer containing 20 mM Tris-HCl (pH 8.0), 150 mM NaCl, 0.5 mM EDTA, 1 mM ATP, 5 mM $MgCl_2$, and 1 mM DTT. Every aliquot taken from the KaiA/KaiB/KaiC-SV mixture incubated at 30°C was used for size-exclusion chromatography analysis at room temperature. A Superdex 200 Increase 10/300 GL (Cytiva) column was connected to a right-angle light scattering (RALS) system (Viscotek TDA305, Malvern) to estimate the molecular masses of the eluted peaks.

### Supplementary Text

Selection of non-phosphorylatable amino acids upon designing phospho-mimicking mutants

Phosphorylatable amino acids, such as serine and threonine, are often substituted with alanine to investigate the functional effects of those phosphoryl modifications. A S431A/T432A KaiC double mutant (KaiC-AA) was used as a phospho-mimic mutant of KaiC-ST (*21*). However, as reported previously (*40*), alanine substitutions at both or either of the dual phosphorylation sites (S431A or T432A) resulted in much higher ATPase activity (25.7 ATP $d^{-1}$ for KaiC-AA) than that observed for fully dephosphorylated KaiC-ST (13.6 ± 1.4 ATP $d^{-1}$) (fig. S2C). By contrast, KaiC-SV is designed to reduce these unwanted side effects by selecting valine instead of alanine because valine has the closest volume (fig. S2A) and topology (fig. S2B) to threonine. KaiC-SV, which has a helical PSw (upper left panel of **Fig. 2**) and similar ATPase activity (13.3 ± 0.7 ATP $d^{-1}$) to KaiC-ST (fig. S2C), exhibits temperature-compensated ATPase- (**Fig. 4B**) and mono–P-cycles (**Fig. 4A**) that are in sharp contrast to the arrhythmicity observed for KaiC-SA and KaiC-AT (fig. S2C). KaiC-CT was designed using the same logic described above to generate a mutant with ATPase activity (9.9 ± 0.7 ATP $d^{-1}$) that does not exceed that of KaiC-ST but also exhibits arrhythmicity (**Fig. 4A**). The present results clearly indicate that S431 is the primary residue responsible for C1–C2 allostery affecting rhythmicity and that special care must sometimes be taken when replacing phosphorylatable amino acids with non-phosphorylatable ones.

Assembly and disassembly dynamics for the KaiA/KaiB/KaiC-pSV ternary complex

After reconstructing the *in vitro* P-cycle for KaiC-SV (fig. S3A), every aliquot taken at a time interval of 13 h was used for size-exclusion chromatography and RALS analyses. When maximally phosphorylated (fig. S3B), KaiC-pSV weakly binds to KaiB. The KaiA/KaiB/KaiC-pSV ternary complex (*8, 9*) begins to form at a dephosphorylating state (fig. S3C), and its accumulation peaks at the maximally dephosphorylated state (fig. S3D). The populated ternary complex is disassembled at a phosphorylating state (fig. S3E). KaiC-SV exhibits minimal allostery to achieve system-level synchronization via KaiA sequestration (*22, 24*) and to drive the assembly–disassembly cycle of the Kai proteins.

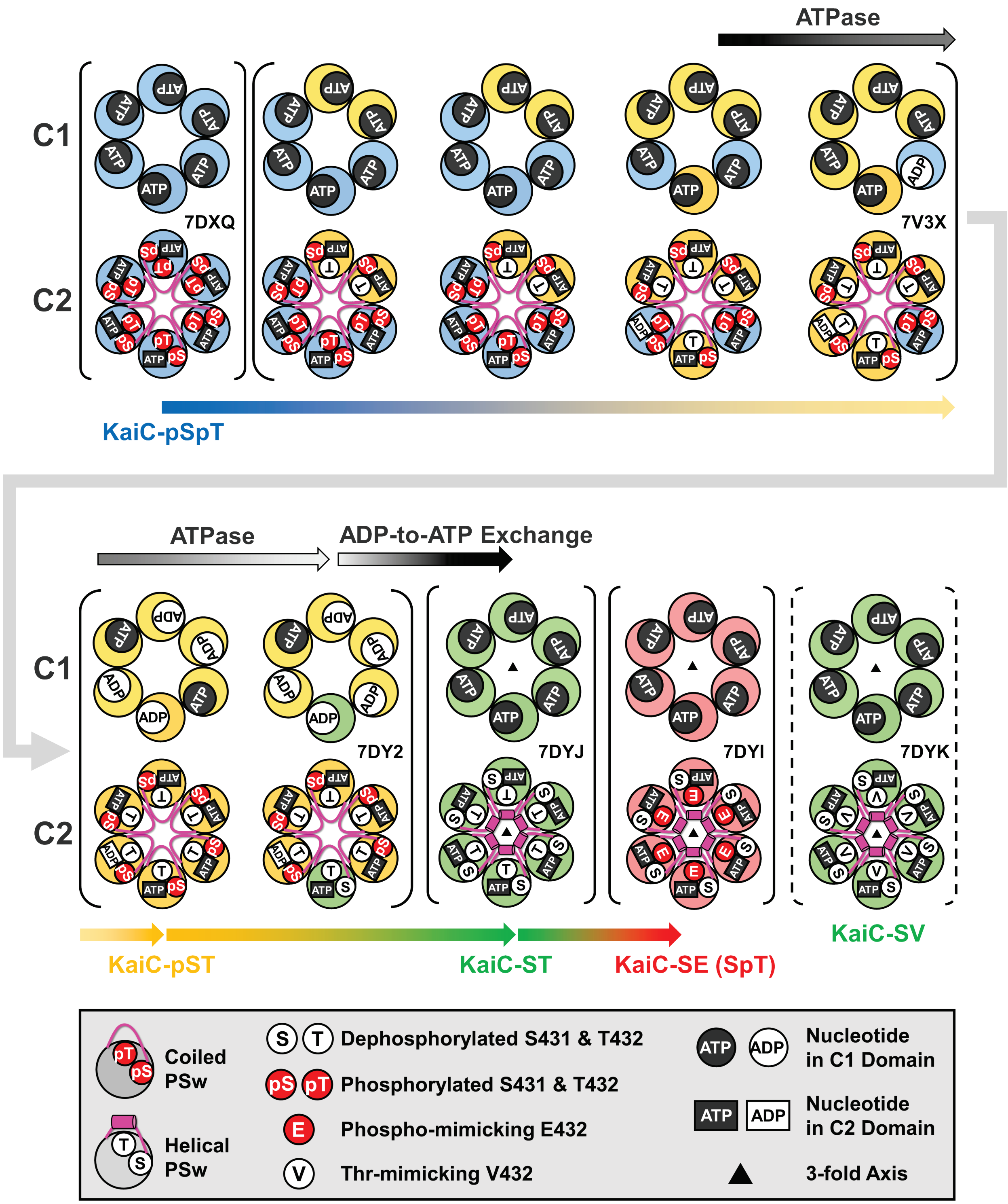


**fig. S1. Schematic drawing of the crystal structure library.** Five crystallographic data sets (in solid parentheses) are aligned in the following order: 7DXQ (a hexamer of KaiC-pSpT in $P2_12_12_1$), 7V3X (four hexamers of partial KaiC-pSpT and KaiC-pST in $P2_1$), 7DY2 (two hexamers of KaiC-pST and partial KaiC-ST in $P2_1$), 7DYJ (a hexamer of KaiC-ST reconstructed in $P6_3$), and 7DYI (a hexamer of KaiC-SE reconstructed in $P6_3$) with annotations in the box below the alignment.

KaiC-SV, a mono–P-cycle oscillator, is coded as 7DYK and drawn as a hexamer according to the $P6_3$ space group in dashed parentheses. The filled triangles in 7DYJ, 7DYI, and 7DYK indicate the crystallographic three-fold axes present at the centers of the hexamers. Arrows above and below the hexameric rings represent biochemical events in the C1 domain and phosphorylation-state transitions in the C2 domain, respectively.

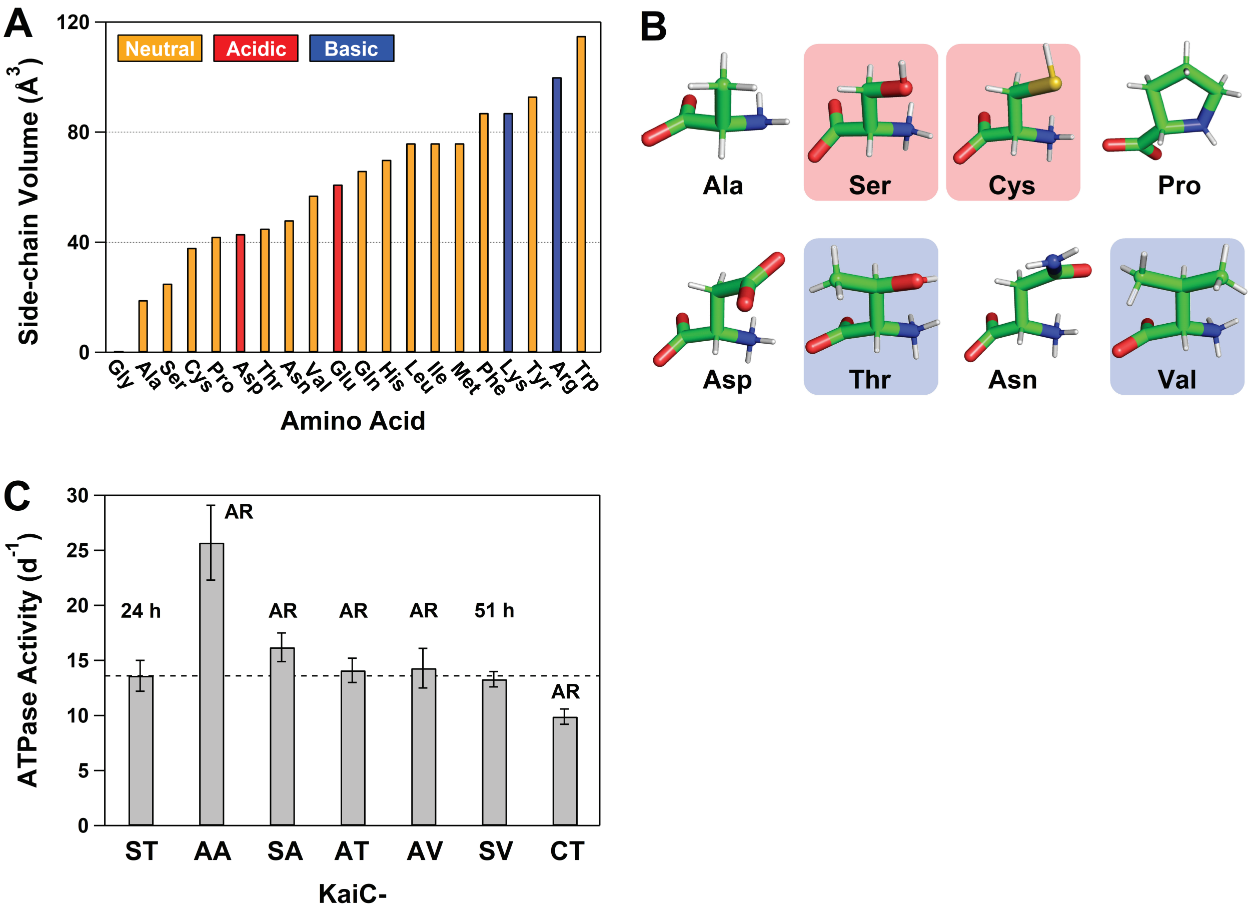


**fig. S2. Candidate amino acid substitutions at dual phosphorylation sites.** (**A**) Side-chain volume and (**B**) topology of amino acid residues. (**C**) Steady-state ATPase activity at 30°C. Values and AR above the error bars indicate the period length or arrhythmicity, respectively, for the *in vitro* P-cycle in the presence of KaiA and KaiB at 30°C.

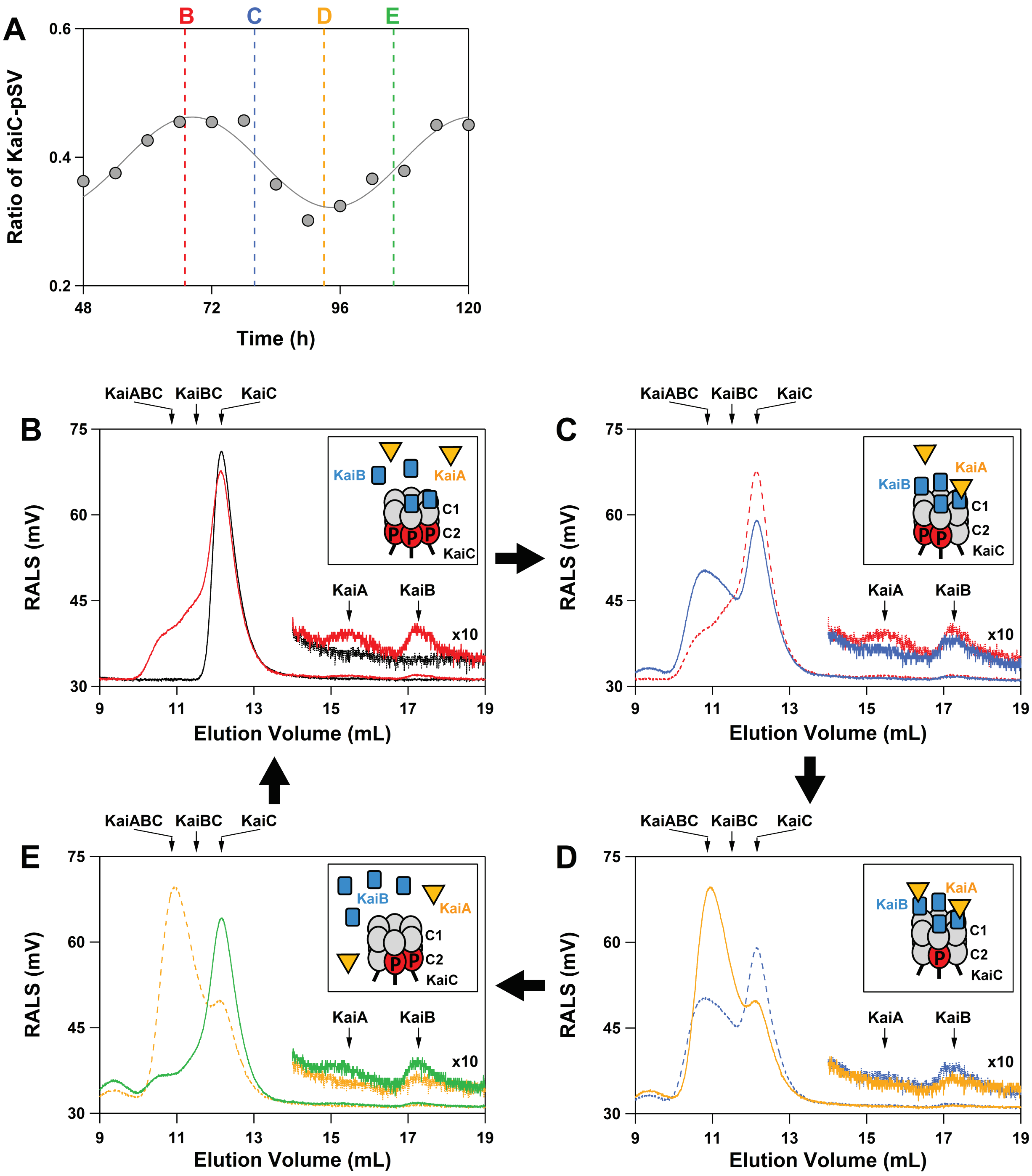


**fig. S3. Assembly and disassembly cycle of the KaiA/KaiB/KaiC-SV ternary complex.** (**A**) P-cycle for KaiC-SV at 30°C. (**B**) Red, (**C**) blue, (**D**) orange, and (**E**) green solid lines correspond to the elution curves for aliquots taken at 67, 80, 93, and 106 h, respectively. Molecular masses of proteins in the eluted fractions were analyzed using right-angle light scattering (RALS). The black solid line depicts the elution curve for KaiC-SV alone. For reference, elution curves for the aliquots taken at 67, 80, and 93 h are shown using red, blue, and orange dotted lines, respectively. The insets correspond to schematic illustrations of the assembled and disassembled states of the ternary complex.

### Table S1. Data collection and refinement statistics.

| **Protein** | KaiC-pSpT | KaiC-pSpT&pST | KaiC-pST | KaiC-ST | KaiC-SE | KaiC-SV |
|---|---|---|---|---|---|---|
| **Data Collection** | | | | | | |
| Space group | $P2_12_12_1$ | $P2_1$ | $P2_1$ | $P6_3$ | $P6_3$ | $P6_3$ |
| Unit cell parameters | | | | | | |
| *a, b, c* (Å) | 92.4, 159.9, 207.24 | 185.5, 205.8, 186.2 | 92.8, 206.6, 168.4 | 94.6, 94.6, 179.9 | 94.3, 94.3, 180.6 | 94.9, 94.9, 180.6 |
| α, β, γ (º) | 90, 90, 90 | 90, 115.1, 90 | 90, 94.7, 90 | 90, 90, 120 | 90, 90, 120 | 90, 90, 120 |
| Wavelength (Å) | 0.9 | 0.9 | 0.9 | 0.9 | 0.9 | 0.9 |
| Resolution range (Å)[a] | 30-2.80 | 50-3.10 | 50-3.04 | 30-2.40 | 48.5-2.64 | 48.5-3.00 |
| | (2.90-2.80) | (3.15-3.10) | (3.15-3.04) | (2.44-2.40) | (2.77-2.64) | (3.11-3.00) |
| Total reflections | 561605 | 883423 | 457569 | 507210 | 280978 | 191896 |
| Unique reflections | 75955 (7451) | 228464 (11426) | 120308 (12016) | 35578 (1775) | 26712 (3551) | 19313 (1788) |
| Redundancy | 7.4 (7.3) | 3.9 (3.8) | 3.8 (3.8) | 14.3 (14.1) | 10.5 (10.4) | 10.4 (10.8) |
| Completeness (%) | 99.9 (100) | 99.9 (100) | 99.9 (100) | 100 (99.8) | 99.9 (99.9) | 99.7 (98.2) |
| $R_{merge}$ (%)[b] | 9.2 (>100) | 10.2 (74.4) | 13.3 (89.9) | 27.0 (>100) | 10.9 (>100) | 13.8 (96.2) |
| (I)/sigma(I) | 21.4 (2.0) | 13.8 (2.1) | 15.6 (2.0) | 36.4 (2.7) | 13.2 (2.1) | 12.9 (2.8) |
| **Model building** | | | | | | |
| Molecular replacement | 2GBL | 2GBL | 2GBL | 2GBL | 2GBL | 7DYJ |
| Total atoms | 21020 | 78739 | 39277 | 6837 | 6669 | 6705 |
| Protien | 20538 | 77135 | 38547 | 6653 | 6510 | 6564 |
| Ligands | 384 | 1512 | 712 | 128 | 128 | 128 |
| Water | 98 | 92 | 18 | 56 | 31 | 13 |
| $R_{work}$ (%)[c] | 25.0 | 27.5 | 26.3 | 26.7 | 29.4 | 27.4 |
| $R_{free}$ (%)[c] | 31.8 | 34.0 | 32.6 | 31.0 | 33.1 | 32.8 |
| R.M.S.D. from ideality | | | | | | |
| Bond length (Å) | 0.002 | 0.006 | 0.002 | 0.002 | 0.002 | 0.002 |
| Bond angles (º) | 1.2 | 1.5 | 1.2 | 1.25 | 1.2 | 1.2 |
| Average B factors (Å$^2$) | 45.7 | 58.0 | 47.0 | 58.4 | 70.2 | 72.7 |
| Rmachandran plot | | | | | | |
| Most favored (%) | 82.6 | 92.2 | 82.3 | 86.9 | 84.5 | 84.2 |
| Allowed (%) | 16.8 | 6.3 | 17.5 | 12.8 | 15.6 | 15.6 |
| Disallowed (%) | 0.6 | 1.4 | 0.2 | 0.2 | 0.0 | 0.1 |
| **PDB code** | 7DXQ | 7V3X | 7DY2 | 7DYJ | 7DYI | 7DYK |

[a]Values in parentheses are for the highest-resolution shell.
[b]$R_{merge} = \Sigma|I-<I>|/\Sigma I$, where I corresponds to the observed intensity of reflections.
[c]$R_{work, free} = \Sigma|F_{obs}|-|F_{calc}|/\Sigma|F_{obs}|$. $R_{free}$ is the cross-validation of the R-factor using the test reflections, 5% of the data, not included in the refinements.
